# Systematic Analysis of KRAS-ligand Interaction Modes and Flexibilities Reveals the Binding Characteristics

**DOI:** 10.1101/2023.02.01.526731

**Authors:** Zheng Zhao, Niraja Bohidar, Philip E. Bourne

**Affiliations:** School of Data Science, University of Virginia, Charlottesville, Virginia 22904, United States of America; Department of Biomedical Engineering, University of Virginia, Charlottesville, Virginia 22904, United States of America

## Abstract

KRAS, a common human oncogene, has been recognized as a critical drug target in treating multiple cancers. After four decades of effort, one allosteric KRAS drug (Sotorasib) has been approved, inspiring more KRAS-targeted drug research. Here we provide the features of KRAS binding pockets and ligand-binding characteristics of KRAS complexes using a structural systems pharmacology approach. Three distinct binding sites (conserved nucleotide-binding site, shallow Switch-I/II pocket, and allosteric Switch-II/α3 pocket) are characterized. Ligand-binding features are determined based on encoded KRAS-inhibitor interaction fingerprints. Finally, the flexibility of the three distinct binding sites to accommodate different potential ligands, based on MD simulation, is discussed. Collectively, these findings are intended to facilitate rational KRAS drug design.

## 1. Introduction

The RAS (Rat sarcoma virus) protein family consists of over 150 members^1^ and belongs to a class of small GTPase enzymes (EC No: 3.6.5.2)^2^. With guanosine nucleotide exchange factors and GTPase-activating proteins, RAS plays a critical role in cell signaling pathways that regulate cell growth, proliferation, differentiation, and apoptosis^3^. Mutations in RAS can impact the preferences of different protein effectors to interact leading to changes in downstream signaling pathways^4^. For example, the G12D-mutated KRAS activates MEK/ERK and PI3K/AKT signaling pathways in the non-small-cell lung cancer (NSCLC) cell line^5^. Indeed, mutations in and overexpression of RAS are found in about 30% of all human cancers, including pancreatic cancer, NSCLC, and breast cancer^6, 7^. Three members of the RAS family, KRAS, HRAS and NRAS, are the most common human oncogenes. Consequently, over the last four decades, attention has been paid to RAS proteins as targets for rational inhibitors design ^8^.

RAS proteins consist of 188 or 189 amino acids, respectively and hence have approximately the same molecular mass (21 kDa) among members and isoforms. RAS proteins contain two domains: the catalytic domain (G domain, amino acids,1-166) and the hypervariable region (amino acids: 167-188/189). The hypervariable regions have less than 20% sequence homology, but retain a highly conserved CAAX motif in the C-terminal region^9^. In contrast, the G domain is highly conserved (Figure 1), comprising a P-loop (residues 10-17), Switch-I region (residues 30-38), Switch-II region (residues 60-76), and Allosteric lobe (residues 87-166). The conformational change of RAS and therefore biological properties are induced by binding GTP or GDP into the nucleotide-binding pocket. Specifically, GTP- and GDP-bound complex structures correspond to active and inactive conformations (Figure 1), respectively^10^. Hence, designing GTP-competitive inhibitors is regarded as one promising approach to cancer treatment. However, targeting RAS is challenging. First, GTP, binds RAS with picomolar affinity^8^, requiring stronger competitive inhibitors to be developed^11^. Second, GTP- or GDP-binding to RAS does not result in substantial conformational changes to form well-defined binding sites. Third, the valley conformation between the Switch regions is so shallow that it is challenging to bind appropriately. Nonetheless, recent progress has been made^12, 13^. For example, Gray’s group developed the covalent catalytic site inhibitors^14, 15^.

**Figure 1.**
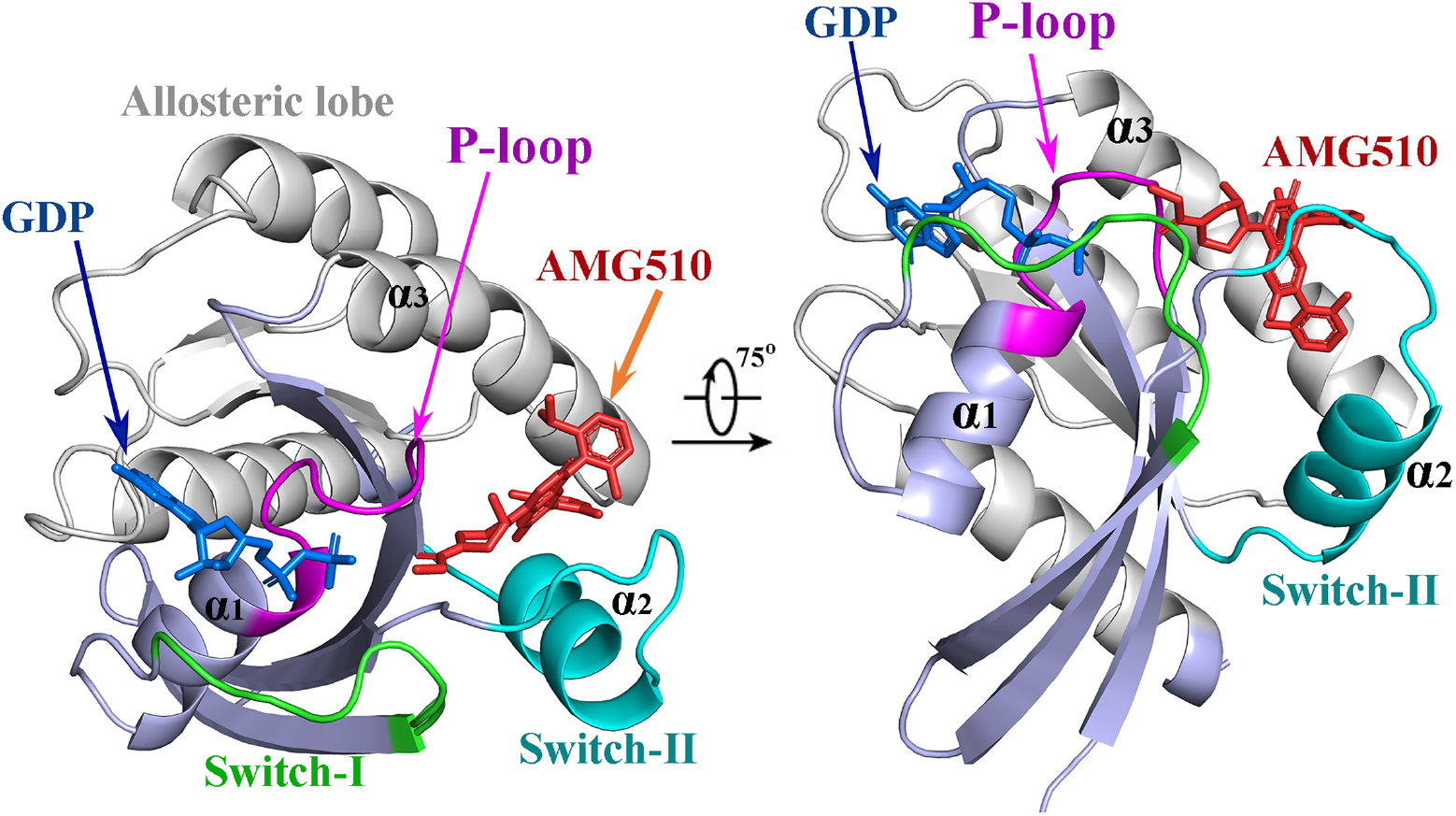
GDP- and AMG510-bound KRAS structure (PDB id: 6oim) marked with GDP (blue), AMG510 (orange), P-loop (purple), Switch-I (green), Switch-II (cyan), Allosteric lobe (gray), comprising helix α1, α2, and α3, respectively.

There’s been direct (KRAS) and indirect (SOS, SHP2) ways to effect the pathway^16^. To date, most RAS-inhibitor development efforts have been focused on targeting KRAS, the most frequently mutated oncogene in human cancers^17^. KRAS has a mutation rate of 95% in human pancreatic cancer, 45% in colorectal cancer, and 45% in lung cancer^18^. Recently progress has been made in designing small molecule inhibitors that target KRAS. On May 28, 2021, the U.S. Food and Drug Administration (FDA) approved the first targeted drug, Sotorasib (previously called AMG510, Figure 1), for adult NSCLC patients, whose genotype shows the KRAS G12C mutation and who have received at least one prior systemic anticancer therapy^10, 19^. Further KRAS G12C inhibitors are in development and dozens are in clinical trials^12, 20^, such as MRTX849, LY3499446, GDC-6036, and JNJ-74699157^21^. To further facilitate KRAS-targeted drug discovery, we conducted a systematic analysis of KRAS binding sites, KRAS-ligand binding characteristics, structural flexibility of KRAS inhibitors, bioactivity, and resistance.

Increasingly available KRAS Protein Data Bank (PDB) experimentally determined structures provide us with an opportunity to holistically navigate KRAS 3D structure space, especially binding sites, using a structural systems pharmacology approach. We first collected all released KRAS structures from the PDB and the dynamic KRAS conformations from KRAS-ligand-bound MD trajectories. From the resultant KRAS PDB structural dataset, we focus on the KRAS-ligand interaction features for all potential and allosteric binding sites by encoding the protein-ligand interaction fingerprints. We then explored the conformational space of ligand-bound KRAS from MD simulations in a simulated physiological environment to uncover the protein-ligand essential interactions and the collective motions for use in prospective drug discovery. Finally, we determined the structural KRAS features of bioactive compounds that inhibit KRAS and KRAS binding sites, hence contributing to KRAS drug design, activity, resistance, and mechanistic understanding.

## 2. Results

We first present the results for the features of KRAS-ligand interactions obtained using the KRAS structural dataset and a function-site interaction fingerprints method^22–24^, with emphasis on every type of binding site. Then we present results on the flexibility of each binding site using an MD simulation method^25^. We compare the dynamic conformational space to that from the KRAS structural (static) dataset. Finally, we highlight details of allosteric inhibitor design for binding into the Switch-II/α3 binding site.

### 2.1. KRAS ligand-binding characteristics

We encoded the ligand-binding characteristics of every KRAS-ligand complex into an interaction-fingerprint binary string (See Method section). A total of 256 interaction fingerprint strings were obtained based on a structural KRAS dataset with 256 crystal KRAS-ligand complexes. Then the 256 interaction-fingerprint strings were aligned based on the secondary structures of the binding sites for all complexes (See Method section) (Table S3). Finally, the 256 interaction-fingerprints were clustered by the hierarchical cluster method (See Method) into three classes: *Class-I, Class-II*, and *Class-III*, possessing distinct binding modes (Figure 2). Each class reveals which specific residues are involved and the nature of the distinct binding modes. For example, ligands Y9Z- and 6ZD-binding complexes (PDB 4nmm and 5kyk, respectively) were clustered into *Class-I* due to their similar binding modes through interacting with common residues, Cys12, Gla15, Lys16, and Ser17. In total, 68 residues (Figure 2, x-axis) are involved in at least one ligand-bound complex in the KRAS dataset. Once classified, these 68 residues constitute distinct binding pockets as discussed subsequently.

**Figure 2.**
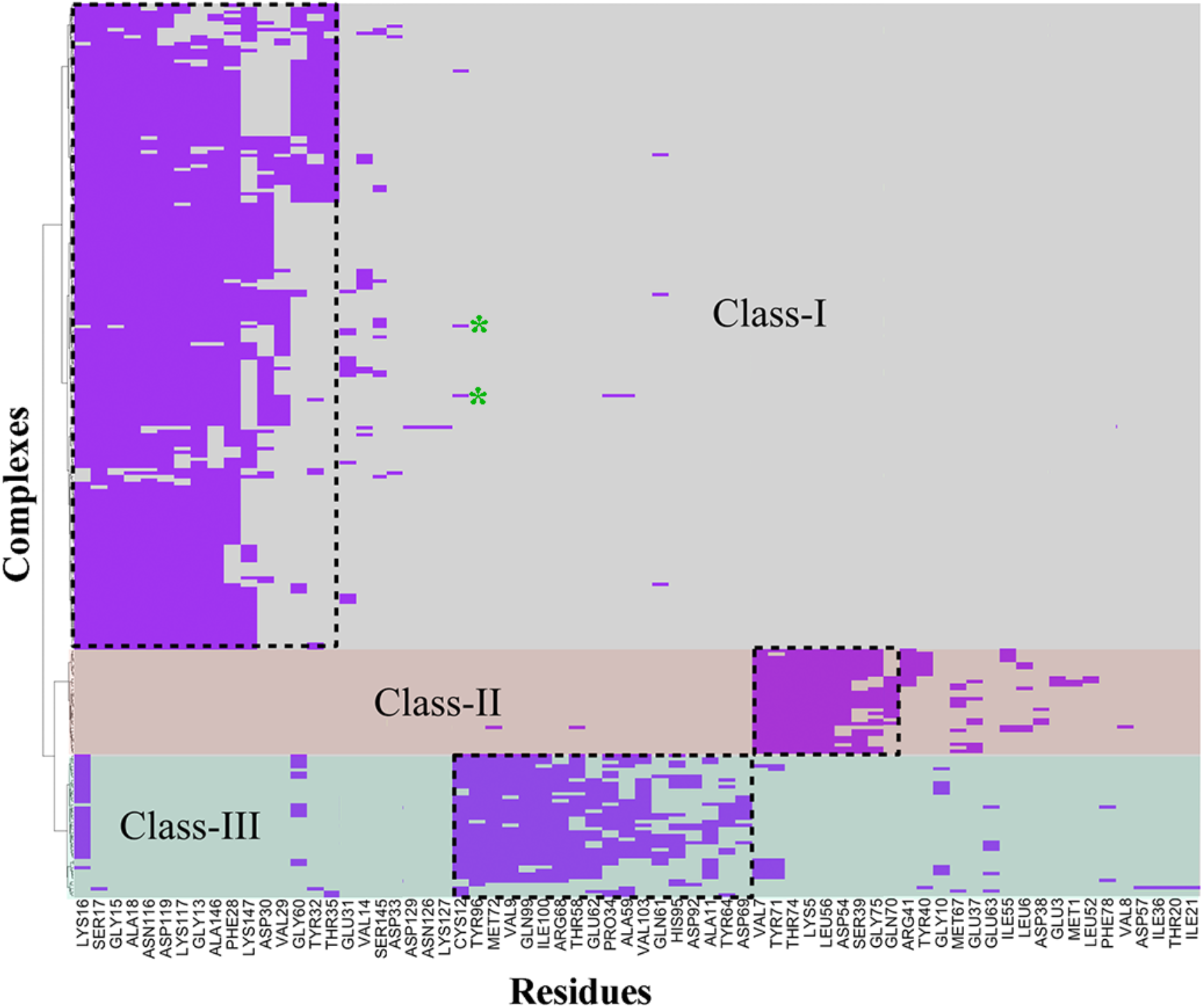
The heatmap represents the binary fingerprint matrix of residue-ligand interactions across the whole structural KRAS-ligand data set. Every row along the y-axis represents one ligand-binding complex. The x-axis represents the residues that constitute the binding pockets. If an amino acid residue has at least one type of interaction with the corresponding ligand, for example, one hydrophobic interaction, the residue in the specific KRAS complex is labeled “1” (purple). Otherwise, the residues are labeled “0” (blank). The interaction-fingerprint strings clustered into three distinct classes: Class-I (gray background), II (light pink background), and III (pale cyan background). The dashes illustrate conserved binding characteristics for each class of binding site. Ligands Y9Z and 6ZD from PDBs 4nmm and 5kyk, respectively, were marked with asterisks (green).

#### Class-I

This category is the largest of the three, consisting of 185 KRAS complexes (Table S4). Residues Gly13, Gla15, Lys16, Ser17, Ala18, Lys117, Phe28, Asp119, Ala146, and Lys147 provide the conserved interactions with all the ligands in this class of KRAS complex (Figure 2). By contrast, the amino acids Cys12, Val14, Val29, Asp30, Glu31, Tyr32, Asp33, Thr35, Gly60, Gln61, Asn116, Lys117, and Ser145 do not occur so frequently (Figure 2). Without conserved interactions these amino acids imply ligands with unusual binding modes and hence selectivity and possibly stronger ligand-binding affinity. For example, ligands Y9Z and 6ZD achieved stronger binding affinity (see below) by uniquely interacting with Cys12 in the G12C-mutated KRAS structures (Figure 3)^26, 27^. Structurally, these amino acids are located in the regions of the P-loop, Switch-I, Switch-II, and Allosteric lobe, respectively, forming the *Class-I* binding pocket (Figure 3a). This binding pocket is also called the “nucleotide-binding site”, where the intrinsic ligands GTP/GDP bind with high affinity.

**Figure 3.**
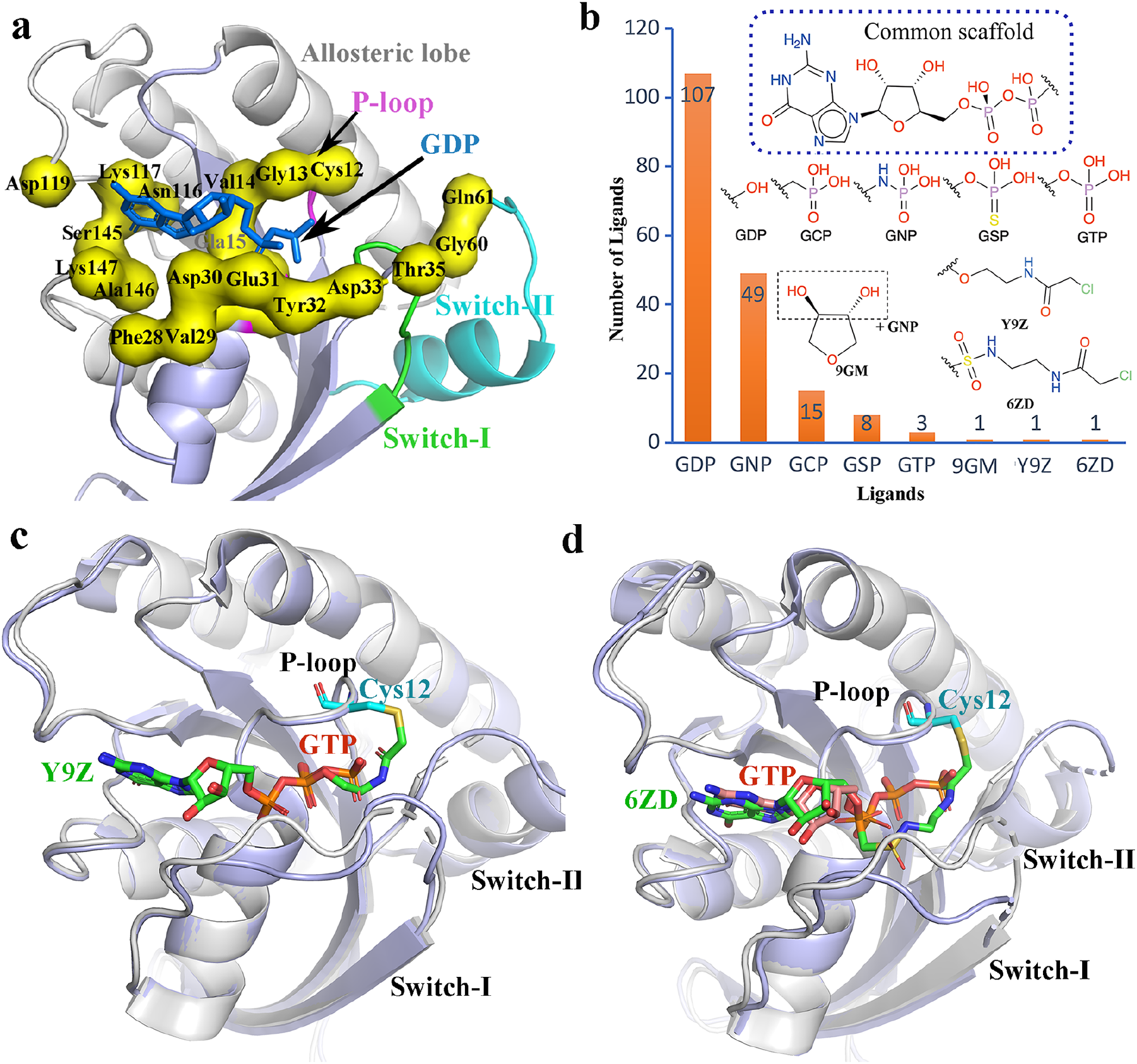
(a) *Class-I* binding site (PDB id 6oim). (b). Distribution of ligands and their 2D structures. (c) Binding mode of Y9Z (green, PDB id 4nmm in light blue) compared with that of GTP (orange, PDB id 5vpi in gray). (d) Comparison of the binding modes of 6ZD (green, PDB id 5kyk in light blue) and GTP (orange, PDB id 5vpi in gray).

Indeed, all ligands studied are highly similar to GTP/GDP and include GCP, GNP, GSP, 9GM, Y9Z, and 6ZD with a common scaffold (Figure 3b). In comparison to GTP, the ligands GCP, GNP and GSP are GTP’s analogs and differ by only one atom. Ligand GDP does not have the functional γ-phosphate. The ligand 9GM has two modifications when compared to GTP: (i) the oxygen atom between the β- and γ-phosphate is a nitrogen atom; (ii) the ribose group has a different ring conformation (Figure 3b). The ligand Y9Z possesses the same nucleotide fragment as GTP, but the γ-phosphate group of GTP is replaced with an electrophilic fragment 2-chloro-N-ethylacetamide, yet occupies the same binding cavity as the γ-phosphate group of GTP. The electrophilic fragment forms the covalent interaction with Cys12 (Figure 3c). Thus, Y9Z has a stronger binding potential than GTP due to the covalent interaction^27^. Ligand 6ZD is also similar to GTP because of the same nucleotide fragment. The β- and γ-phosphates of GTP are replaced by chloroacetamide. Chloroacetamide is an electrophilic fragment and also has a covalent interaction with Cys12 (*Ki=* 9 nM, Figure 3d). In summary, the *Class-I* ligands bind to the nucleotide-binding site and share the same core structure (i.e., nucleotide) to block the pro-oncogenic RAS/MAPK signaling pathway. This suggests that this nucleotide-binding site is conserved to accommodate GTP/GDP or its analogs. Additionally, in the nucleotide-binding pocket, establishing a covalent warhead to GTP analogs is a potential way to design GTP-competitive inhibitors for treating cancers with the K12C genotype mutation.

#### Class-II

There are 30 KRAS-ligand-binding complexes in this class (Figure 2, Table S4). The interaction fingerprints are predominantly from residues Lys5, Val7, Ser39, Asp54, Leu56, Gly70, Tyr71, Thr74, and Gly75, which are located in the regions of the β1-3, and Switch-II (Figure 4a) and form the core structure of this binding site. Near the core structure, more vicinal residues, including Leu6, Glu37, Asp38, Tyr40, Arg41, Ile55, and Met67, also provide specific interactions to accommodate the diverse ligands. Collectively, these residues form the *Class-II* binding site, called Switch-I/II (Figure 4a). The Switch-I/II binding site does not have any overlap with the *Class-I* nucleotide-binding site (Figure 3a). From the 30 complexes in this class, the 30 ligands form three distinct clusters (Figure 4b) when using a hierarchical clustering method (see Method section). Cluster1 ligands interact predominantly with the core structure of the *Class-II* binding pocket^28^ but also extend to interact with the residues on the β3 strand. For example, the ligand F8Q not only has one hydrogen-bond interaction with Thr74 on helix α2, but also accommodates a hydrophobic groove mainly formed by the Ser39-Tyr-40-Arg41 peptide on the β2 strand (Figure 4c). Cluster2 ligands also have binding interactions with the core residues of the *Class-II* binding site^29^. Compared to Cluster1 ligands, Cluster2 ligands have interactions with the residues near Switch-II. (Figure 4d). One example from Cluster2 is ligand 0QV, which also has a hydrogenbond interaction with Thr74 on helix α2 as does the ligand F8Q in Cluster1. Ligand 0QV extends into the Switch-II area and accommodates a polar, solvent-exposed sub-pocket mainly formed by Glu37 in the Switch-I loop and Glu62 in the Switch-II loop. Cluster3 ligands not only bind to the center of the *Class-II* binding site, but also extend into the N-terminal region of β1. For example, one end of the R6W ligand (Figure 4e) binds to the core part of the Switch-I/II pocket^30^. The other end, located at the surface, close to the N-terminal of the β1, influences the protein-protein interaction with another domain of the dimer. In total, Cluster1-3 ligands in *Class-II* all bind to the core part of the *Class-II* binding site, but each cluster extends in different directions to achieve the desired binding affinity by interacting with vicinal residues (Figure 4c-e). The *Class-II* binding site is not a typical pocket, but rather a shallow valley (Figure 4a), which requires one side of the ligand to strongly interact with β1-3, Switch-I, and Switch-II, raising the possibility that the other side of the ligand interacts with other proteins. Recently, Kessler et al. showed that the small molecule BI-2852, binds to the *Class-II* binding site with nanomolar affinity, inhibiting KRAS-effector interactions^30^. Conversely, Timothy et al. provided an alternative explanation with BI-2852 inducing a KRAS dimerization to achieve inhibitory activity^31^.

**Figure 4.**
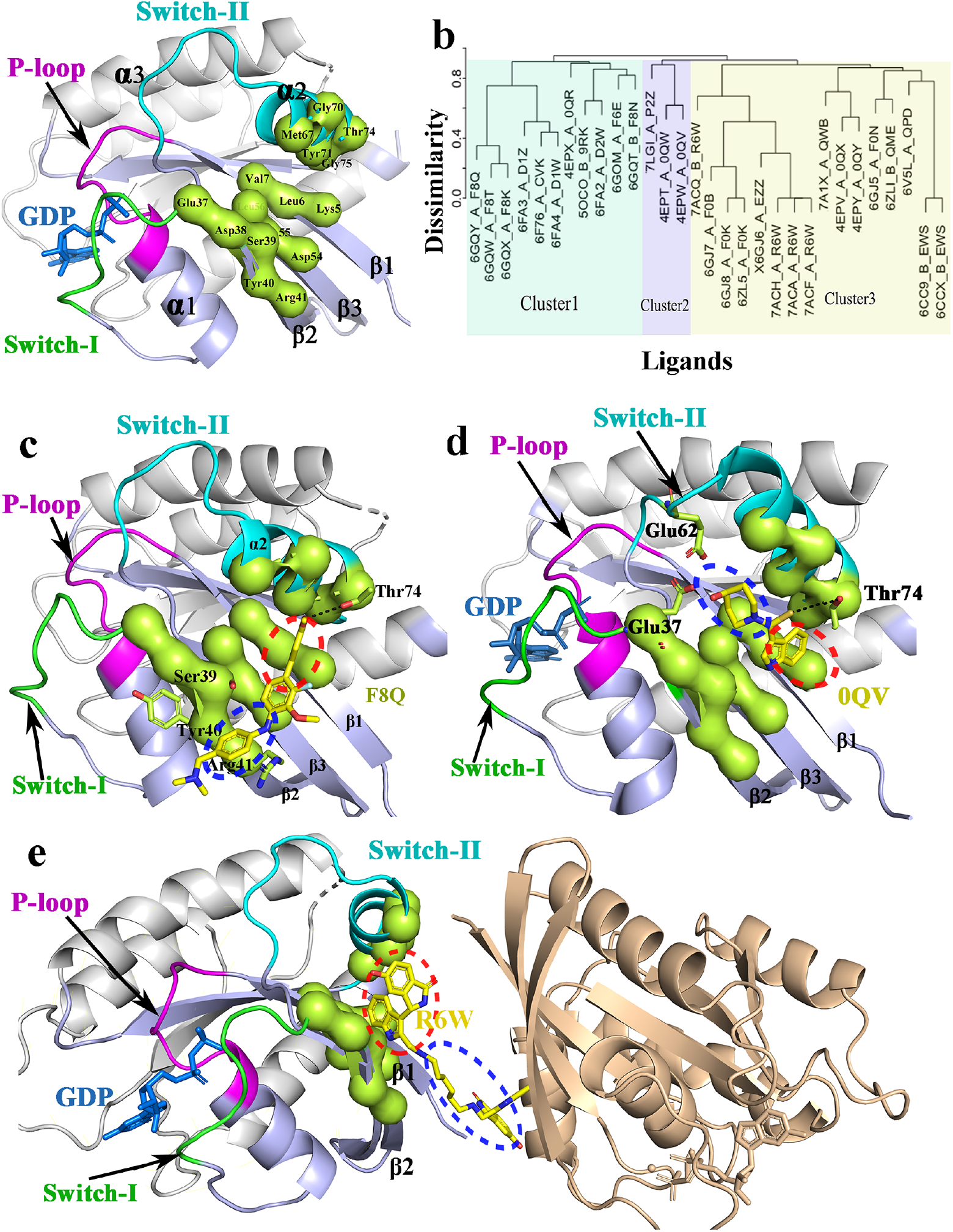
(a) *Class-II* binding site (PDB id 6oim). (b) Hierarchical cluster analysis of ligands binding into the *Class-II* binding pocket. (c) Cluster1 binding mode as shown in PDB 6gqy with ligand F8Q (yellow). (d) Cluster2 binding mode as shown in PDB 4epw with ligand 0QV (yellow) (e) Cluster3 binding mode as shown in PDB 7acf with ligand R6W (yellow). The red dashed ellipses show the ligands interacting with the core residues of the binding pocket. The blue dashed ellipses show the ligands interacting with the extended residues of the binding pocket in each cluster.

#### Class-III

There are 41 KRAS-ligand complexes in this class. The binding site, here called Switch-II/α3, is comprised of 18 residues: Val9, Ala11, Cys12, Pro34, Thr58, Ala59, Gln61, Glu62, Tyr64, Arg68, Asp69, Met72, Asp92, His95, Tyr96, Gln99, Ile100, andVal103. Only Val9, Ala11, and Cys12 are located on the P-loop and one residue, Pro34, is located at the Switch-I region. The other 14 residues are located within the Switch-II loop and helix α3 (Figure 5a). Clustering of the ligands reveals three clusters (A, B, and C), representing three different binding modes, including the substantial allosteric effect of this binding site (Figure 5b). In Cluster-A, the ligand QY5 binds into one end of the *Class-III* binding site between the P-loop, Switch-I, and Switch-II, and has no interactions with helix α3 (Figure 5c)^32^. Indeed, Switch-I deviates away from helix α3 (Figure 3c). In Cluster-B, Switch-II deviates away from helix α3 to accommodate the Cluster-B ligands. For example, a Cluster-B ligand, 21S, is located between the P-loop and Switch-II, and does not have any interactions with helix α3 (Figure 5d)^20^. Compared to Clusters-A and - B, Cluster-C ligands induce a bigger occupying space and fully interact with the P-loop, Switch-I, Switch-II, and helix α3 (Figure 5a). The ligand AMG510, an approved drug^10^, binds to this induced allosteric pocket. Additionally, the function group of acrylamide forms a covalent interaction with Cys12 (Figure 5a) to permanently lock the G12C KRAS in an inactive state^19^. In sum, this binding site flexibly accommodates different ligands, presenting distinct allosteric effects for each ligand.

**Figure 5.**
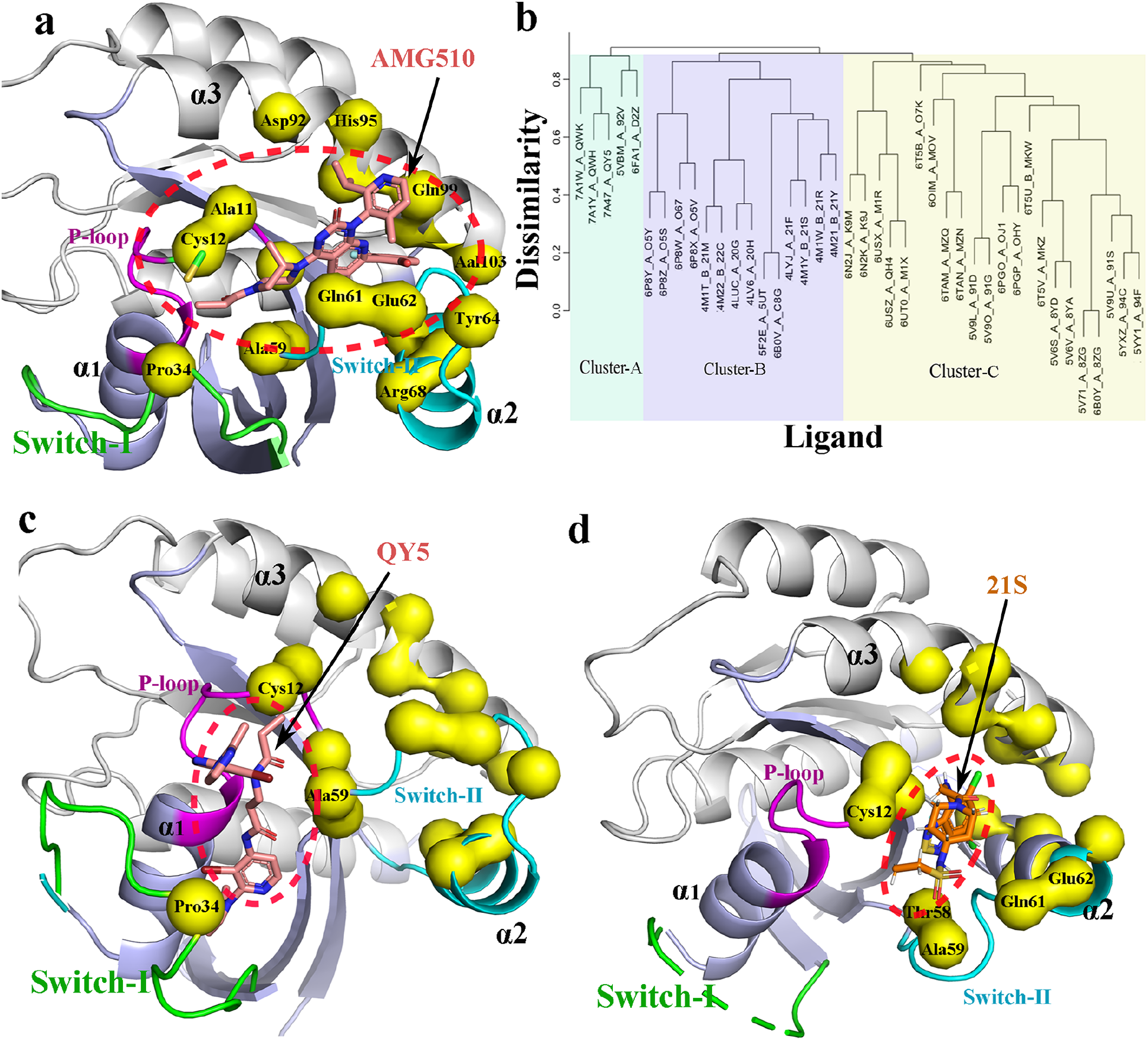
(a) *Class-III* KRAS binding site (PDB id: 6oim) and the binding mode of Cluster-C ligands. (b) Clustered ligands in the *Class-III* binding site. (c) The binding mode of the Cluster-A ligands (PDB id 7a47). (d) The binding mode of the Cluster-B ligands (PDB id:4m1y). The red dashed ellipses mark the binding zones of the *Class-III* ligands.

### 2.2. Flexibility of KRAS binding sites

To explore the flexibility of KRAS binding sites and their relationships to ligand binding, we examined three different KRAS systems (namely, apo KRAS, GDP-bound KRAS, and AMG510-bound KRAS) using 360 ns molecular dynamics simulations (See Methods, Figure S1-2).

The volume of the nucleotide-binding sites are very similar (medium = ~ 235 Å^3^, Figure 6a) in the AMG510- and GDP-bound systems, indicating that the nucleotide-binding site does not undergo a substantial change upon either binding a ligand (AMG510) at the Switch-II/α3 binding site or binding a GDP ligand. The bound ligands form stable amino acid-ligand interactions resulting in fewer flexibility changes in the P-loop and Switch-I loop at the nucleotide-binding site. The less flexible GDP-bound pocket is similar to that in the crystal structures (median = ~205 Å^3^, Figure 6b). This is in agreement with the aforementioned discussion that the ligands inside the nucleotide-binding site are similar (Figure 3b). Thus, unsurprisingly, thus far the currently reported KRAS inhibitors binding to this nucleotide-binding site are GDP’s analogs^12^, such as SML-8-73-1, SML-10-70-1, and XY-02-075^14, 15, 27^. The AMG510 ligand, binding into the Switch-II/α3 binding site, forms a covalent interaction with Cys12 of the P-loop, which means that AMG510 impacts the nucleotide-binding site to adopt the same conformation as the GDP-binding pocket conformation. From this viewpoint, it will be feasible to design inhibitors that bind to the Switch-II/α3 binding site in the GDP-bound KRAS conformation^10, 19^. In the apo KRAS system, the nucleotide-binding site does exhibit obvious flexibility with volume changes from 50 to 250 Å^3^(median = ~197 Å^3^). However, the largest volume (about 250 Å^3^) of the nucleotide-binding site is similar to that in the GDP- or AMG510-bound systems (Figure 6a). Thus, the nucleotide-binding site is conserved and prefers to bind GDP and its analogs (Figure 3b).

**Figure 6.**
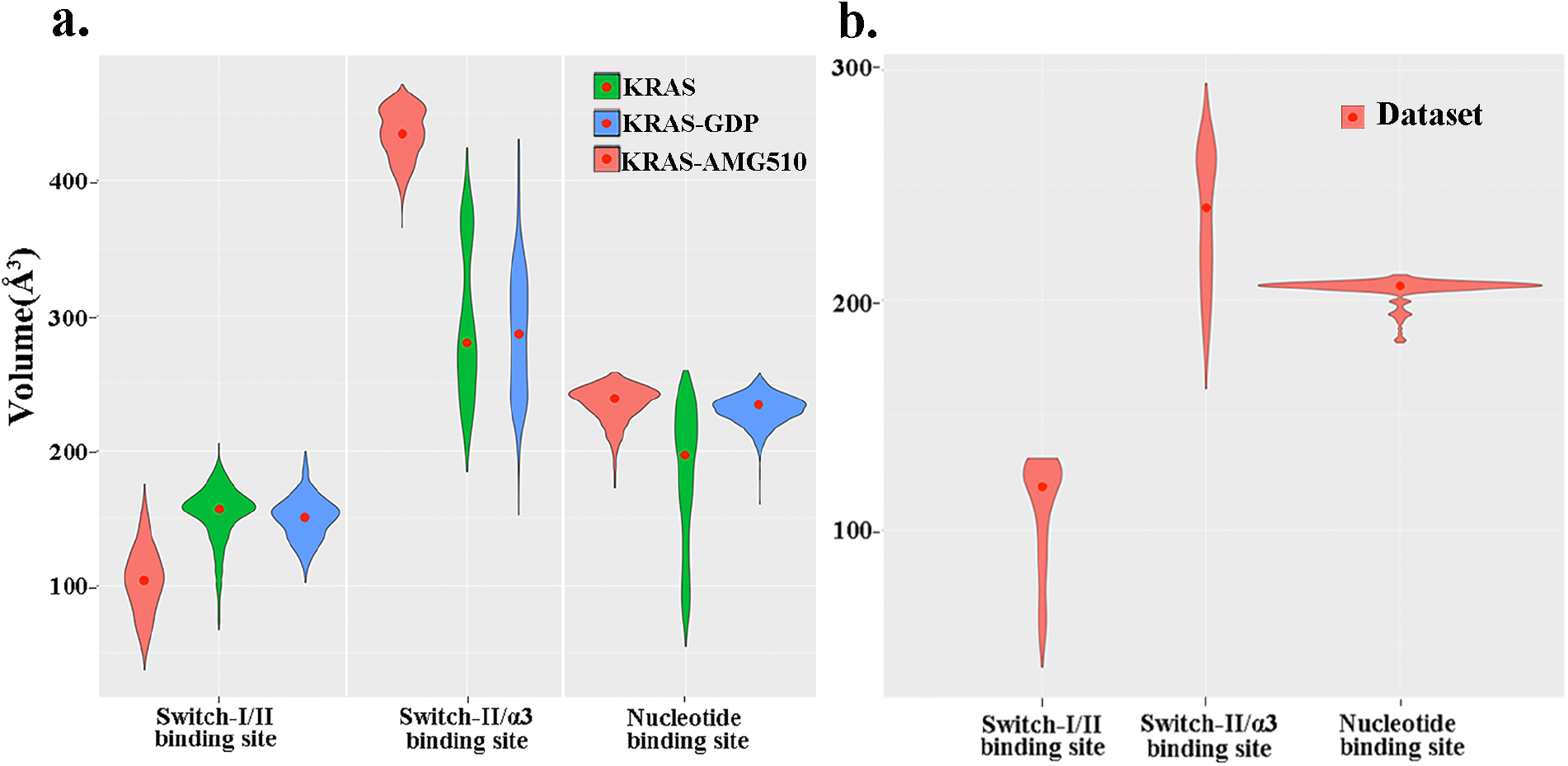
The flexibility of KRAS binding sites. (a-b) Volumatic analysis of KRAS binding sites from the conformation space from MD simulations (a) and KRAS PDB dataset (b), respectively. The width of the violin box is proportional to the frequency in volume of the binding pocket. The red dot in the violin box represents the median.

Out of the three classes of KRAS binding pocket, the Switch-I/II binding site is the smallest since it is very shallow and on the surface of KRAS domain. In the apo KRAS system, the median size of the Switch-I/II binding site is ~ 155 Å^3^. Compared to the apo KRAS system, this Switch-I/II binding site does not exhibit a big conformational change (median = ~150 Å^3^) when GDP is bound to the nucleotide binding site. In contrast, when AMG510 is bound to the Switch-II/α3 binding pocket, the Switch-I/II binding site becomes smaller (median = ~ 100 Å^3^), The Switch-II segment (residues 60-76) shows flexibility with a large dynamic magnitude (Figure S2). Moreover, the Switch-II’s motion is towards the Switch-I/II binding site to accommodate AMG510 in the Switch-II/α3 binding site.

Of the three types of binding pocket, the Switch-II/α3 binding site is the largest with the medians ~280, ~285, and ~430 Å^3^ in apo KRAS, GDP-bound, and AMG510-bound MD systems, respectively. Notably, the size of the Switch-II/α3 binding site shows the largest fluctuations ranging from ~150 Å^3^ to ~430 Å^3^ when it is not occupied (apo KRAS or GDP-binding KRAS), which suggests that the Switch-II/α3 binding site provides substantial flexibility to aid in allosteric drug discovery. When AMG510 occupies the Switch-II/α3 binding pocket, the size of the binding pocket becomes much larger (median: ~430 Å^3^) and an induced binding mode is formed. Interestingly, in crystal structures (Figure 6b), this binding site also exhibited the same tendency for flexibility although not as much (median: ~235 Å^3^) as that in the MD simulations, which reveal a larger conformational change (Figure 6a) for the allosteric Switch-II/α3 binding site under physiological condition. Important information is designing more promising KRAS inhibitors. FDA approval of AMG510 shows the potential of the binding site by utilizing its flexibility to design allosteric inhibitors^19^. Taken together, the three pockets have variable volumes and exhibit substantial flexibility, important features in the design of KRAS inhibitors.

### 3. Conclusion

We have systematically analyzed KRAS binding pockets and the binding characteristics of the corresponding ligands known to bind to KRAS PDB structures. From the analysis of ligandbinding features, the binding pockets can be classified into three classes called the nucleotide-binding site, the Switch-I/II pocket, and the Switch-II/α3 pocket^12, 19^. The nucleotide-binding site is a conserved binding pocket, where GTP and analogs containing a fragment of nucleotide prefer to bind. The tail of the GTP molecule (i.e., γ-phosphate) is close to the P-loop and Switch-II, and could be replaced using highly electrophilic groups to design GTP-competitive inhibitors, such as the inhibitor 6ZD (PDB id 5kyk), which forms a covalent interaction with the P-loop by mimicking the β- and γ-phosphates of GTP^14, 15, 27^. We also found that the Switch-I/II binding site is a shallow pocket located at the surface of the KRAS domain. This shallow pocket mediates a possible protein-protein interaction, such as found in the ligand R6W which is located between two KRAS domains (PDB id 7acf, Figure 4e). The third class of KRAS pocket is the Switch-II/α3 pocket, and shows larger flexibility (Figure 6b). Upon binding different ligands, the Switch-I region, or the Switch-II region always undergo significant change to accommodate the specific ligands (Figure 5a, c-d), demonstrating a potential allosteric effect and providing a more diverse target potential in the design and discovery of KRAS inhibitors.

We explored the dynamic flexibilities of the binding pockets upon ligand binding. The GDP-binding system shows this nucleotide-binding site is relatively conserved. The AMG510-binding KRAS system shows that the Switch-II/α3 pocket has the largest induced fluctuation where ligands are bound. Encouragingly, in the apo KRAS systems, the binding sites all present a larger range of fluctuation (Figure 6a). Likewise, in the AMG510-bound system, the nucleotide-binding pocket also shows a substantial fluctuation akin to the GDP-binding KRAS system (Figure 6), which implies the potential to dynamically screen for novel allosteric KRAS inhibitors in vitro or in vivo using the apo KRAS system or GDP-bound system. Taken together, these findings provide insights into the design and the characteristics of KRAS inhibitors.

### 4. Methods

In this study, we applied a structural systems pharmacology pipeline (Figure 7) by combining the function-site interaction fingerprint method and the volumetric analysis of binding sites to reveal KRAS pocket characteristics for drug discovery.

**Figure 7.**
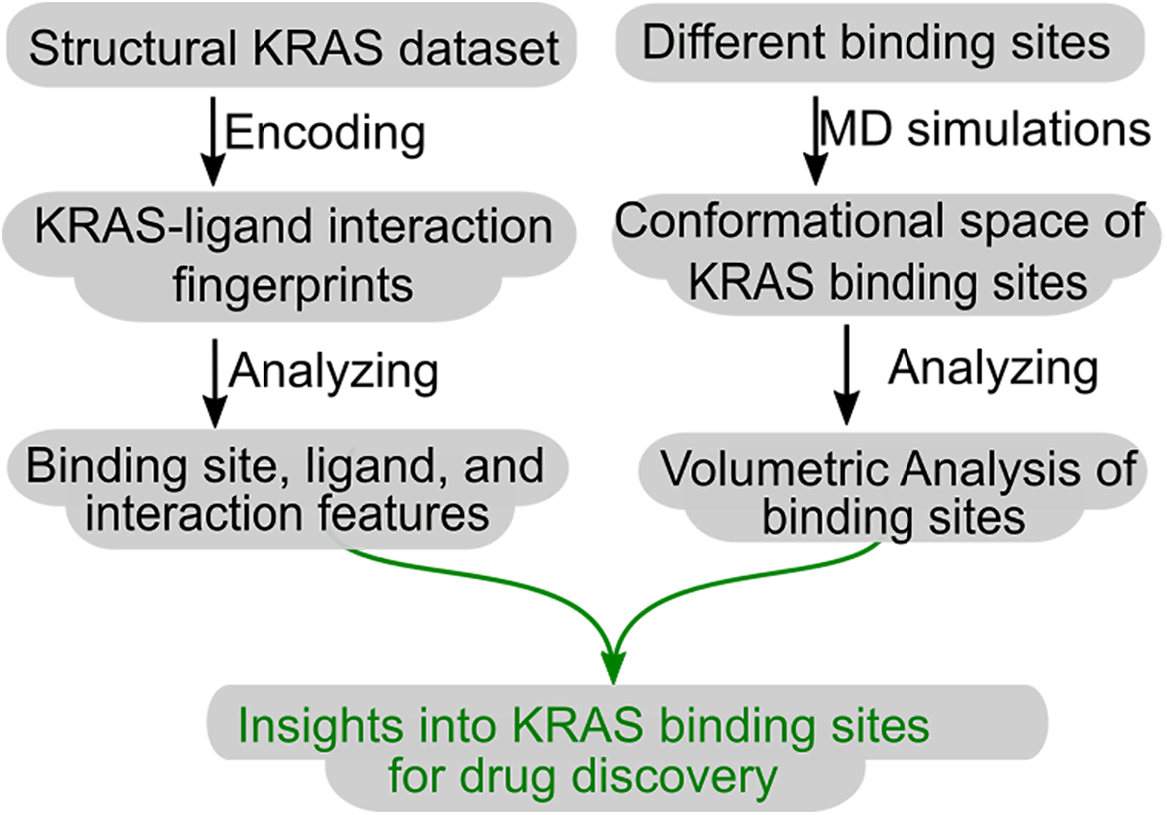
The structural systems pharmacology scheme used in this study.

### 4.1. Structural ligand-bound KRAS dataset

We downloaded all the KRAS PDB structures from the PDB database (https://www.rcsb.org)^22^ based on the list of PDB identifiers associated with the UniProt entry P01116 from The Universal Protein Resource (UniProt)^33^ as of October 16^th^, 2021. In total, 206 PDB structures were collected including 194 X-ray structures, 1 Cryo-EM structure, and 11 NMR structures (Table S1). To build upon this ligand-bound KRAS dataset, we removed small molecules, such as metal ions, polyatomic anions, or any other intermediate molecules. Then we filtered out 15 partial KRAS structures (sequence length < 100 amino acids) and 6 structures without ligands. The final curated dataset comprised 185 KRAS complexes containing 256 ligands (Table S2).

### 4.2 Encoding protein-ligand interaction fingerprints

We used the Function-site Interaction Fingerprint (Fs-IFP) method to encode the protein-ligand interaction features into binary strings^23, 24^. The Fs-IFP method has been extensively applied to explore ligand binding characteristics, virtual screening, and drug mechanisms at a proteome scale^34–37^. Briefly, the method comprises three steps. Step 1 aligns all protein-ligand complexes from the given dataset using a sequence-independent 3D binding sites aligning method, such as MultiBind^38^, mTM-align^39^, or SMAP^40^. Step 2 encodes the interaction fingerprints for each protein-ligand complex. Every residue interacting with the ligand is encoded as a 7-bit substring, which represents 7 types of interaction^41^, including electrostatic interactions, π-stack interactions, and hydrophobic interactions. Step 3 combines the aligned amino acid sequences with the encoded interaction fingerprints to obtain comparable interaction fingerprints.

In this paper, we performed the alignment of multiple binding sites and encoding interaction fingerprints using the SMAP^40^ and IChem^42^ software, respectively. The hierarchical cluster analysis of all of the interaction fingerprint strings was carried out using Ward’s minimum variance method^43^ in the R package: RStudio (https://www.rstudio.com).

### 4.3 Ligand similarity

We used the ScreenMD function add-in in ChemAxon (version 22.7.0) to calculate ligand similarity^44^. First, every ligand was represented as a 1024-bit extended-connectivity fingerprint (ECFP), which is a type of circular topological fingerprints designated for collating molecular features, similarities, and structure-activity relationships^45^. Then the pairwise similarity was calculated using the Tanimoto coefficient, i.e.,

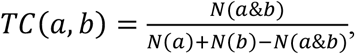

in which, a and b are two equal-length binary fingerprints, & is the and-operator for the bitwise comparison, and *N*(*x*) is the number of 1 bits given any binary fingerprint *x*. Finally, the Tanimoto dissimilarity matrix was obtained from all the pairwise similarities and was clustered using hierarchical cluster analysis from the complete linkage method available in the R package: RStudio (https://www.rstudio.com).

### 4.4. MD simulations

The apo-, GDP-bound-, and AMG510-bound-KRAS structures were obtained from the Protein Data Bank with PDB ids: 4ldj, 5v9u, and 6oim, respectively^22^. The repeated Chain B in 5v9u and the crystal water molecules in all structures were removed. Missing loops were added using the Modeller software^46^. Then, the force field parameters and the topology files were constructed using the online ParamChem server^47^ for small molecules GDP and AMG510. Correspondingly, the CHARMM36 force field^48^ was used for the protein. Then, every system was solved in a rectangular box filled with TIP3P water molecules. The side lengths of the rectangular box are 81.16, 74.62, and 80.53 Å, respectively, which ensure an 18 Å buffer from any solute atom. The protonation of all of the charged amino acids was determined based on a pH of 7.0. Finally, the counterions (Na^+^ and Cl^-^) were added to make sure the system had an ion concentration of 0.20 M and electroneutrality.

First, the three KRAS systems were optimized using a normal restrained protocol: 500 steps of energy minimization with a Ca restraint constant of 10 kcal.mol^-1^.Å^-2^; 2ns MD simulations with a Ca restraint constant of 5.0 kcal.mol^-1^.Å^-2^; 2ns MD simulations with a Ca restraint constant of 2.0 kcal.mol^-1^.Å^-2^. Then, one 120 ns MD simulation was carried out under standard conditions (298.15 K, 1 atm) for each system and the last 100 ns of the trajectories were used to analyze the flexibility of the binding sites. The MD simulation software ACEMD was used for all MD simulations^25^. The particle-mesh Ewald (PME) summation was applied with a cutoff distance of 9.0 Å for van der Waals and electrostatic interactions. The temperature and pressure were kept using the Langevin thermostat and Monte Carlo barostat, respectively^25^. The integration time step is 4fs with all bond constraints and repartition hydrogen mass to 4.0 au.

The volumetric size of each class of binding site for every conformation was calculated using the volumetric analysis of surface properties (VASP) software^49^, in which all computations were carried out at 0.5 Å resolution. Finally, the change of every class of binding sites was shown with a violin plot.

## 5. Data and Software Availability

The KRAS structure data in the paper were downloaded from the Protein Data Bank, located at https://www.rcsb.org. The software encoding protein-ligand interaction fingerprints is IChem from Dr. Didier Rognan’s structural chemogenomics group (http://bioinfo-pharma.u-strasbg.fr/labwebsite/download.html). The license can be requested from Dr. Didier Rognan for academic use. All the MD simulations were performed using the commercial software ACEMD, located at https://www.acellera.com/acemd/. The ACEMD toolbox running in a single GPU is free for all. The ligand similarity in this paper was calculated using the ScreenMD addin from the commercial software ChemAxon (https://chemaxon.com). This company offers different licenses, such as academic teaching licenses, and some free applications for compound property calculations (https://chemicalize.com/welcome). The binding pocket volume size calculations were performed using VASP, which is open-source software under the Gnu General Public License v3.0 and can be downloaded at http://www.cse.lehigh.edu/~chen/index.htm. The secondary structural alignments were performed using the software SMAP, which can be freely downloaded (http://compsci.hunter.cuny.edu/~leixie/smap/smap.html) for academic users.

## 6. Supporting Information

The released KRAS PDB structures (Table S1); The structural ligand-bound KRAS dataset includes 185 KRAS complexes containing 256 ligands (Table S2). The complete list for 256 interaction-fingerprint strings (Table S3, XLSX). The list of clustered complexes for each class (Table S4). Ca-RMSDs of different trajectories for all three systems (Figure S1). Ca-RMSFs of different trajectories for all three systems (Figure S2).

## 7. Acknowledgment

We acknowledge Research Computing at The University of Virginia and at the Extreme Science and Engineering Discovery Environment (XSEDE) (BIO210030, Z.Z.) for providing computational resources and technical support that have contributed to the results reported within this publication. This work described here was supported by the University of Virginia (P.E.B).

