## Supplementary material for "Systematic Analysis of KRAS-ligand Interaction Modes and Flexibilities Reveals the Binding Characteristics": Table 1,2, and 4 and Figure 1-2

**Table S1.** The released KRAS PDB structures.

| PDB ID | METHOD | RESOLUTION | SEQUENCE | PDB ID | METHOD | RESOLUTION | SEQUENCE |
|---|---|---|---|---|---|---|---|
| **1D8D** | X-ray | 2 | 178-188 | **6EPP** | X-ray | 2.4 | 1-169 |
| **1D8E** | X-ray | 3 | 178-188 | **6F76** | X-ray | 2.2 | 1-169 |
| **1KZO** | X-ray | 2.2 | 169-173 | **6FA1** | X-ray | 1.97 | 1-167 |
| **1KZP** | X-ray | 2.1 | 169-173 | **B/E** | 1-169 | | |
| **1N4P** | X-ray | 2.65 | 185-189 | **6FA2** | X-ray | 2.6 | 1-167 |
| **1N4Q** | X-ray | 2.4 | 185-189 | **6FA3** | X-ray | 1.82 | 1-167 |
| **1N4R** | X-ray | 2.8 | 185-189 | **6FA4** | X-ray | 2.02 | 1-169 |
| **1N4S** | X-ray | 2.6 | 185-189 | **6GJ5** | X-ray | 1.5 | 1-169 |
| **3GFT** | X-ray | 2.27 | 1-164 | **6GJ6** | X-ray | 1.76 | 1-169 |
| **4DSN** | X-ray | 2.03 | 2-164 | **6GJ7** | X-ray | 1.67 | 1-169 |
| **4DSO** | X-ray | 1.85 | 2-164 | **6GJ8** | X-ray | 1.65 | 1-167 |
| **4EPR** | X-ray | 2 | 1-164 | **6GOD** | X-ray | 1.71 | 2-173 |
| **4EPT** | X-ray | 2 | 1-164 | **6GOE** | X-ray | 1.6 | 2-169 |
| **4EPV** | X-ray | 1.35 | 1-164 | **6GOF** | X-ray | 1.98 | 2-173 |
| **4EPW** | X-ray | 1.7 | 1-164 | **6GOG** | X-ray | 2.05 | 1-169 |
| **4EPX** | X-ray | 1.76 | 1-164 | **6GOM** | X-ray | 1.63 | 1-167 |
| **4EPY** | X-ray | 1.8 | 1-164 | **6GQT** | X-ray | 1.69 | 1-167 |
| **4L8G** | X-ray | 1.52 | 1-169 | **6GQW** | X-ray | 2.8 | 1-167 |
| **4LDJ** | X-ray | 1.15 | 1-164 | **6GQX** | X-ray | 2.2 | 1-167 |
| **4LPK** | X-ray | 1.5 | 1-169 | **6GQY** | X-ray | 2.75 | 1-167 |
| **4LRW** | X-ray | 2.15 | 1-169 | **6H46** | X-ray | 2.22 | 1-166 |
| **4LUC** | X-ray | 1.29 | 1-169 | **6H47** | X-ray | 1.7 | 1-166 |
| **4LV6** | X-ray | 1.5 | 1-169 | **6JTN** | X-ray | 1.9 | 19-Oct |
| **4LYF** | X-ray | 1.57 | 1-169 | **6JTO** | X-ray | 1.7 | 19-Oct |
| **4LYH** | X-ray | 1.37 | 1-169 | **6JTP** | X-ray | 1.9 | 18-Oct |
| **4LYJ** | X-ray | 1.93 | 1-169 | **6M9W** | X-ray | 1.5 | 2-169 |
| **4M1O** | X-ray | 1.57 | 1-169 | **6MBQ** | X-ray | 1.35 | 2-166 |
| **4M1S** | X-ray | 1.55 | 1-169 | **6MBT** | X-ray | 1.45 | 1-169 |
| **4M1T** | X-ray | 1.7 | 1-169 | **6MBU** | X-ray | 1.45 | 1-169 |
| **4M1W** | X-ray | 1.58 | 1-169 | **6MNX** | X-ray | 2.2 | 1-169 |
| **4M1Y** | X-ray | 1.49 | 1-169 | **6MQG** | X-ray | 1.5 | 3-169 |
| **4M21** | X-ray | 1.94 | 1-169 | **6MQN** | X-ray | 1.6 | 1-169 |
| **4M22** | X-ray | 2.09 | 1-169 | **6MS9** | X-ray | 1.49 | 1-169 |
| **4NMM** | X-ray | 1.89 | 1-164 | **6MTA** | X-ray | 2.15 | 1-169 |
| **4OBE** | X-ray | 1.24 | 1-164 | **6N2J** | X-ray | 1.8 | 1-169 |
| **4PZY** | X-ray | 1.88 | 1-164 | **6N2K** | X-ray | 1.72 | 1-169 |
| **4PZZ** | X-ray | 1.4 | 1-164 | **6O36** | X-ray | 2 | 1-167 |
| **4Q01** | X-ray | 1.29 | 1-164 | **6O46** | X-ray | 1.9 | 1-167 |
| **4Q02** | X-ray | 1.7 | 1-164 | **6O4Y** | X-ray | 1.58 | 14-Jul |

| | | | | | | | |
|---|---|---|---|---|---|---|---|
| **4Q03** | X-ray | 1.2 | 1-164 | **6O4Z** | X-ray | 1.5 | 12-May |
| **4QL3** | X-ray | 1.04 | 1-164 | **6O51** | X-ray | 1.55 | 14-Jun |
| **4TQ9** | X-ray | 1.49 | 1-164 | **6O53** | X-ray | 1.4 | 14-May |
| **4TQA** | X-ray | 1.13 | 1-164 | **6OB2** | X-ray | 2.85 | 1-169 |
| **4WA7** | X-ray | 1.99 | 1-164 | **6OB3** | X-ray | 2.1 | 1-169 |
| **5F2E** | X-ray | 1.4 | 1-169 | **6OIM** | X-ray | 1.65 | 1-169 |
| **5KYK** | X-ray | 2.7 | 1-167 | **6P0Z** | X-ray | 1.01 | 2-169 |
| **5MLA** | X-ray | 2.19 | 1-166 | **6P8W** | X-ray | 2.1 | 1-169 |
| **5MLB** | X-ray | 3.22 | 1-166 | **6P8X** | X-ray | 2.11 | 1-169 |
| **5O2S** | X-ray | 3.22 | 1-166 | **6P8Y** | X-ray | 2.31 | 1-169 |
| **5O2T** | X-ray | 2.19 | 1-166 | **6P8Z** | X-ray | 1.65 | 1-169 |
| **5OCG** | X-ray | 1.48 | 2-189 | **6PGO** | X-ray | 1.6 | 1-169 |
| **5OCO** | X-ray | 1.66 | 1-169 | **6PGP** | X-ray | 1.5 | 1-169 |
| **5-Oct** | X-ray | 2.07 | 1-169 | **6PQ3** | X-ray | 1.75 | 1-169 |
| **5TAR** | X-ray | 1.9 | 2-164 | **6PTS** | NMR | - | 1-186 |
| **5TB5** | X-ray | 2 | 2-164 | **6PTW** | NMR | - | 1-186 |
| **5UFE** | X-ray | 2.3 | 1-166 | **6QUU** | X-ray | 1.48 | 1-169 |
| **5UFQ** | X-ray | 2.2 | 1-166 | **6QUV** | X-ray | 1.48 | 1-169 |
| **5UK9** | X-ray | 1.89 | 1-166 | **6QUW** | X-ray | 1.24 | 1-169 |
| **5UQW** | X-ray | 1.5 | 1-164 | **6QUX** | X-ray | 1.62 | 1-169 |
| **5US4** | X-ray | 1.83 | 1-164 | **6T5B** | X-ray | 1.37 | 1-169 |
| **5USJ** | X-ray | 1.94 | 1-164 | **6T5U** | X-ray | 1.72 | 1-166 |
| **5V6S** | X-ray | 1.7 | 1-169 | **6T5V** | X-ray | 1.31 | 1-169 |
| **5V6V** | X-ray | 1.72 | 1-169 | **6TAM** | X-ray | 1.64 | 1-169 |
| **5V71** | X-ray | 2.23 | 1-167 | **6TAN** | X-ray | 1.16 | 1-169 |
| **5V9L** | X-ray | 1.98 | 1-167 | **6USX** | X-ray | 2.27 | 1-169 |
| **5V9O** | X-ray | 1.56 | 1-167 | **6USZ** | X-ray | 2.03 | 1-169 |
| **5V9U** | X-ray | 1.38 | 1-169 | **6UT0** | X-ray | 1.94 | 1-169 |
| **5VBM** | X-ray | 1.49 | 1-169 | **6V5L** | NMR | - | 1-169 |
| **5VP7** | X-ray | 1.7 | 1-169 | **6V65** | X-ray | 2.76 | 1-169 |
| **5VPI** | X-ray | 1.62 | 1-169 | **6V6F** | X-ray | 2.54 | 1-169 |
| **5VPY** | X-ray | 2 | 1-169 | **6VC8** | X-ray | 2.5 | 1-169 |
| **5VPZ** | X-ray | 1.85 | 1-169 | **6VJJ** | X-ray | 1.4 | 1-169 |
| **5VQ0** | X-ray | 2.3 | 1-169 | **6W4E** | NMR | - | 2-186 |
| **5VQ1** | X-ray | 1.78 | 1-169 | **6W4F** | NMR | - | 2-186 |
| **5VQ2** | X-ray | 1.96 | 1-169 | **6WGN** | X-ray | 1.6 | 1-169 |
| **5VQ6** | X-ray | 1.99 | 1-169 | **6XGU** | X-ray | 2.7 | 1-169 |
| **5VQ8** | X-ray | 2.3 | 1-169 | **6XGV** | X-ray | 2.11 | 1-169 |
| **5W22** | X-ray | 1.76 | 1-169 | **6XHA** | X-ray | 2.87 | 1-169 |
| **5WHA** | X-ray | 2.04 | 1-166 | **6XHB** | X-ray | 2.5 | 1-169 |
| **5WHB** | X-ray | 2.18 | 1-166 | **6YR8** | X-ray | 1.9 | 1-166 |
| **5WHD** | X-ray | 1.64 | 1-166 | **6YXW** | X-ray | 2.06 | 1-167 |

| | | | | | | | |
|---|---|---|---|---|---|---|---|
| **5WHE** | X-ray | 1.91 | 1-166 | **6ZL5** | X-ray | 1.65 | 1-169 |
| **5WLB** | X-ray | 1.72 | 1-166 | **6ZLI** | X-ray | 1.73 | 1-169 |
| **5WPM** | X-ray | 1.72 | 1-166 | **7A1W** | X-ray | 1.76 | 1-169 |
| **5XCO** | X-ray | 1.25 | 1-169 | **7A1X** | X-ray | 1.32 | 1-164 |
| **5YXZ** | X-ray | 1.7 | 1-169 | **7A1Y** | X-ray | 2 | 1-164 |
| **5YY1** | X-ray | 1.69 | 1-169 | **7A47** | X-ray | 2.16 | 1-169 |
| **6ARK** | X-ray | 1.75 | 1-169 | **7ACA** | X-ray | 1.57 | 1-169 |
| **6ASA** | X-ray | 2.54 | 1-167 | **7ACF** | X-ray | 1.91 | 1-169 |
| **6ASE** | X-ray | 1.55 | 1-169 | **7ACH** | X-ray | 1.9 | 1-169 |
| **6B0V** | X-ray | 1.29 | 1-169 | **7ACQ** | X-ray | 1.86 | 1-169 |
| **6B0Y** | X-ray | 1.43 | 1-169 | **7C40** | X-ray | 2.52 | 1-167 |
| **6BOF** | X-ray | 1.4 | 2-169 | **7C41** | X-ray | 2.28 | 1-167 |
| **6BP1** | X-ray | 2 | 1-169 | **7EYX** | X-ray | 1.82 | 2-169 |
| **6CC9** | NMR | - | 1-186 | **7F0W** | X-ray | 1.39 | 2-169 |
| **6CCH** | NMR | - | 1-186 | **7KFZ** | electron microscopy | 3.47 | 1-169 |
| **6CCX** | NMR | - | 1-186 | **7LC1** | X-ray | 2.35 | 1-169 |
| **6CU6** | X-ray | 1.5 | 1-169 | **7LC2** | X-ray | 2.7 | 1-169 |
| **6E6F** | X-ray | 3.4 | 1-166 | **7LGI** | NMR | - | 1-169 |
| **6E6G** | X-ray | 1.93 | 1-166 | **7NY8** | X-ray | 1.8 | 1-167 |
| **6EPL** | X-ray | 2.55 | 1-169 | **7ROV** | X-ray | 1.32 | 1-189 |
| **6EPM** | X-ray | 2.5 | 1-169 | **7RSC** | NMR | - | 2-186 |
| **6EPN** | X-ray | 2.5 | 1-169 | **7RSE** | NMR | - | 2-186 |
| **6EPO** | X-ray | 2.4 | 1-169 | | | | |

**Table S2.** The struc.tural ligand-bound KRAS dataset including 185 KRAS complexes containing 256 ligands.

| PDB IDS | CHAIN | LIGAND | PDB IDS | CHAIN | LIGAND | PDB IDS | CHAIN | LIGAND |
|---|---|---|---|---|---|---|---|---|
| **3GFT** | A | GNP | **5VQ0** | A | GDP | **6P8X** | A | GDP |
| **4DSN** | A | GCP | **5VQ1** | A | GDP | **6P8X** | A | O5V |
| **4DSO** | A | GSP | **5VQ2** | A | GTP | **6P8Y** | A | GDP |
| **4EPR** | A | GDP | **5VQ6** | A | GSP | **6P8Y** | A | O5Y |
| **4EPT** | A | 0QW | **5VQ8** | A | GDP | **6P8Z** | A | GDP |
| **4EPT** | A | GDP | **5W22** | A | GDP | **6P8Z** | A | O5S |
| **4EPV** | A | 0QX | **5WHA** | A | GDP | **6PGO** | A | GDP |
| **4EPV** | A | GDP | **5WHB** | A | GDP | **6PGO** | A | OJ1 |
| **4EPW** | A | 0QV | **5WHD** | A | GDP | **6PGP** | A | GDP |
| **4EPW** | A | GDP | **5WHE** | A | GNP | **6PGP** | A | OHY |
| **4EPX** | A | 0QR | **5WLB** | A | GNP | **6QUU** | A | GCP |
| **4EPX** | A | GDP | **5WPM** | A | GNP | **6QUV** | A | GCP |
| **4EPY** | A | 0QY | **5XCO** | A | GDP | **6QUW** | A | GCP |
| **4EPY** | A | GDP | **5YXZ** | A | 94C | **6QUX** | A | GCP |
| **4L8G** | A | GDP | **5YXZ** | A | GDP | **6T5B** | A | GDP |
| **4LDJ** | A | GDP | **5YY1** | A | 94F | **6T5B** | A | O7K |
| **4LPK** | A | GDP | **5YY1** | A | GDP | **6T5V** | A | GDP |
| **4LRW** | A | GDP | **6ARK** | A | GDP | **6T5V** | A | MKZ |
| **4LUC** | A | 20G | **6ASA** | A | GDP | **6V5L** | A | GDP |
| **4LUC** | A | GDP | **6ASE** | A | GDP | **6V5L** | A | QPD |
| **4LV6** | A | 20H | **6B0V** | A | C8G | **6V65** | C | GNP |
| **4LV6** | A | GDP | **6B0V** | A | GDP | **6V6F** | C | GNP |
| **4LYF** | A | GDP | **6B0Y** | A | 8ZG | **6VC8** | A | GNP |
| **4LYH** | A | GDP | **6B0Y** | A | GDP | **6VJJ** | A | GNP |
| **4LYJ** | A | 21F | **6BOF** | A | GDP | **6W4E** | B | GSP |
| **4LYJ** | A | GDP | **6BP1** | A | GCP | **6W4F** | B | GDP |
| **4M1O** | A | GDP | **6CC9** | B | EWS | **6WGN** | A | GNP |
| **4M1S** | A | GDP | **6CC9** | B | GNP | **6XGU** | A | GNP |
| **4M1T** | B | 21M | **6CCH** | B | GNP | **6XGV** | A | GNP |
| **4M1T** | B | GDP | **6CCX** | B | EWS | **6XHA** | A | GNP |
| **4M1W** | B | 21R | **6CCX** | B | GNP | **6XHB** | A | GNP |
| **4M1W** | B | GDP | **6CU6** | A | GNP | **6YR8** | A | GDP |
| **4M1Y** | B | 21S | **6E6F** | A | GNP | **6YXW** | A | GDP |
| **4M1Y** | B | GDP | **6E6G** | A | GDP | **6ZL5** | A | GDP |
| **4M21** | B | 21Y | **6F76** | A | CVK | **6ZL5** | A | F0K |
| **4M21** | B | GDP | **6F76** | A | GNP | **6ZLI** | B | GCP |
| **4M22** | B | 22C | **6FA1** | A | D2Z | **6ZLI** | B | QME |
| **4M22** | B | GDP | **6FA1** | A | GNP | **7A1W** | A | GDP |
| **4NMM** | A | Y9Z | **6FA2** | A | D2W | **7A1W** | A | QWK |

| | | | | | | | | |
|---|---|---|---|---|---|---|---|---|
| **4OBE** | A | GDP | **6FA2** | A | GNP | **7A1X** | A | GDP |
| **4PZY** | A | GDP | **6FA3** | A | D1Z | **7A1X** | A | QWB |
| **4PZZ** | A | GDP | **6FA3** | A | GNP | **7A1Y** | A | GDP |
| **4Q01** | A | GDP | **6FA4** | A | D1W | **7A1Y** | A | QWH |
| **4Q02** | A | GDP | **6FA4** | A | GNP | **7A47** | A | GDP |
| **4Q03** | A | GDP | **6GJ5** | A | F0N | **7A47** | A | QY5 |
| **4QL3** | A | GDP | **6GJ5** | A | GCP | **7ACA** | A | GCP |
| **4TQ9** | A | GDP | **6GJ6** | A | EZZ | **7ACA** | A | R6W |
| **4TQA** | A | GDP | **6GJ6** | A | GCP | **7ACF** | A | GCP |
| **4WA7** | A | GDP | **6GJ7** | A | F0B | **7ACF** | A | R6W |
| **5F2E** | A | 5UT | **6GJ7** | A | GCP | **7ACH** | A | GCP |
| **5F2E** | A | GDP | **6GJ8** | A | F0K | **7ACH** | A | R6W |
| **5KYK** | A | 6ZD | **6GJ8** | A | GCP | **7ACQ** | B | GDP |
| **5MLA** | A | GSP | **6GOD** | A | GNP | **7ACQ** | B | R6W |
| **5MLB** | A | GDP | **6GOE** | A | GNP | **7C40** | A | GDP |
| **5O2S** | A | GDP | **6GOF** | A | GNP | **7C41** | A | GDP |
| **5O2T** | A | GSP | **6GOG** | A | GNP | **7EYX** | A | GDP |
| **5OCG** | A | GNP | **6GOM** | A | F6E | **7F0W** | A | GDP |
| **5OCO** | B | 9RK | **6GOM** | A | GNP | **7KFZ** | C | GNP |
| **5OCO** | B | GNP | **6GQT** | B | F8N | **7LC1** | A | GNP |
| **5-Ост** | A | GNP | **6GQT** | B | GNP | **7LC2** | A | GNP |
| **5TAR** | A | GDP | **6GQW** | A | F8T | **7LGI** | A | GDP |
| **5TB5** | A | GDP | **6GQW** | A | GNP | **7LGI** | A | P2Z |
| **5UFE** | A | GNP | **6GQX** | A | F8K | **7NY8** | A | GDP |
| **5UFQ** | A | GNP | **6GQX** | A | GNP | **7ROV** | A | GCP |
| **5UK9** | A | GDP | **6GQY** | A | F8Q | **7RSC** | A | GSP |
| **5UQW** | A | GDP | **6GQY** | A | GNP | **7RSE** | A | GSP |
| **5US4** | A | GDP | **6H46** | A | GDP | **6MNX** | A | GTP |
| **5USJ** | A | GNP | **6M9W** | A | GDP | **6MS9** | A | GDP |
| **5V6S** | A | 8YD | **6MBQ** | A | GNP | **6MTA** | A | GNP |
| **5V6S** | A | GDP | **6MBT** | A | GDP | **6PQ3** | A | GDP |
| **5V6V** | A | 8YA | **6MBU** | A | GDP | **6PTS** | B | GNP |
| **5V6V** | A | GDP | **6MQG** | A | GDP | **6PTW** | B | GNP |
| **5V71** | A | 8ZG | **6MQN** | A | GDP | **6T5U** | B | GDP |
| **5V71** | A | GDP | **6N2J** | A | GDP | **6T5U** | B | MKW |
| **5V9L** | A | 91D | **6N2J** | A | K9M | **6TAM** | A | GDP |
| **5V9L** | A | GDP | **6N2K** | A | GDP | **6TAM** | A | MZQ |
| **5V9O** | A | 91G | **6N2K** | A | K9J | **6TAN** | A | GDP |
| **5V9O** | A | GDP | **6O36** | A | GNP | **6TAN** | A | MZN |
| **5V9U** | A | 91S | **6O46** | A | GNP | **6USX** | A | GDP |
| **5V9U** | A | GDP | **6OB2** | A | GNP | **6USX** | A | M1R |
| **5VBM** | A | 92V | **6OB3** | A | GNP | **6USZ** | A | GDP |

| | | | | | | | | |
|---|---|---|---|---|---|---|---|---|
| **5VBM** | A | GDP | **6OIM** | A | GDP | **6USZ** | A | QH4 |
| **5VP7** | A | GDP | **6OIM** | A | MOV | **6UT0** | A | GDP |
| **5VPI** | A | GTP | **6P0Z** | A | GDP | **6UT0** | A | M1X |
| **5VPY** | A | 9GM | **6P8W** | A | GDP | | | |
| **5VPZ** | A | GSP | **6P8W** | A | O67 | | | |

**Table S4.** The list of clustered complexes for each class.

| CLASSES | PDBID_CHAIN_LIGANDNAME |
|---|---|
| CLASS-I | 3GFT_A_GNP 4DSN_A_GCP 4DSO_A_GSP 4EPR_A_GDP 4EPT_A_GDP 4EPV_A_GDP<br>4EPW_A_GDP 4EPX_A_GDP 4EPY_A_GDP 4L8G_A_GDP 4LDJ_A_GDP 4LPK_A_GDP<br>4LRW_A_GDP 4LUC_A_GDP 4LV6_A_GDP 4LYF_A_GDP 4LYH_A_GDP 4LYJ_A_GDP<br>4M1O_A_GDP 4M1S_A_GDP 4M1T_B_GDP 4M1W_B_GDP 4M1Y_B_GDP 4M21_B_GDP<br>4M22_B_GDP 4NMM_A_Y9Z 4OBE_A_GDP 4PZY_A_GDP 4PZZ_A_GDP 4Q01_A_GDP<br>4Q02_A_GDP 4Q03_A_GDP 4QL3_A_GDP 4TQ9_A_GDP 4TQA_A_GDP 4WA7_A_GDP<br>5F2E_A_GDP 5KYK_A_6ZD 5MLA_A_GSP 5MLB_A_GDP 5O2S_A_GDP 5O2T_A_GSP<br>5OCG_A_GNP 5OCO_B_GNP 5OCT_A_GNP 5TAR_A_GDP 5TB5_A_GDP 5UFE_A_GNP<br>5UFQ_A_GNP 5UK9_A_GDP 5UQW_A_GDP 5US4_A_GDP 5USJ_A_GNP 5V6S_A_GDP<br>5V6V_A_GDP 5V71_A_GDP 5V9L_A_GDP 5V9O_A_GDP 5V9U_A_GDP 5VBM_A_GDP<br>5VP7_A_GDP 5VPI_A_GTP 5VPY_A_9GM 5VPZ_A_GSP 5VQ0_A_GDP 5VQ1_A_GDP<br>5VQ2_A_GTP 5VQ6_A_GSP 5VQ8_A_GDP 5W22_A_GDP 5WHA_A_GDP 5WHB_A_GDP<br>5WHD_A_GDP 5WHE_A_GNP 5WLB_A_GNP 5WPM_A_GNP 5XCO_A_GDP<br>5YXZ_A_GDP 5YY1_A_GDP 6ARK_A_GDP 6ASA_A_GDP 6ASE_A_GDP 6B0V_A_GDP<br>6B0Y_A_GDP 6BOF_A_GDP 6BP1_A_GCP 6CC9_B_GNP 6CCH_B_GNP 6CCX_B_GNP<br>6CU6_A_GNP 6E6F_A_GNP 6E6G_A_GDP 6F76_A_GNP 6FA1_A_GNP 6FA2_A_GNP<br>6FA3_A_GNP 6FA4_A_GNP 6GJ5_A_GCP 6GJ6_A_GCP 6GJ7_A_GCP 6GJ8_A_GCP<br>6GOD_A_GNP 6GOE_A_GNP 6GOF_A_GNP 6GOG_A_GNP 6GOM_A_GNP 6GQT_B_GNP<br>6GQW_A_GNP 6GQX_A_GNP 6GQY_A_GNP 6H46_A_GDP 6M9W_A_GDP 6MBQ_A_GNP<br>6MBT_A_GDP 6MBU_A_GDP 6MNX_A_GTP 6MQG_A_GDP 6MQN_A_GDP<br>6MS9_A_GDP 6MTA_A_GNP 6N2J_A_GDP 6N2K_A_GDP 6O36_A_GNP 6O46_A_GNP<br>6OB2_A_GNP 6OB3_A_GNP 6OIM_A_GDP 6P0Z_A_GDP 6P8W_A_GDP 6P8X_A_GDP<br>6P8Y_A_GDP 6P8Z_A_GDP 6PGO_A_GDP 6PGP_A_GDP 6PQ3_A_GDP 6PTS_B_GNP<br>6PTW_B_GNP 6QUU_A_GCP 6QUV_A_GCP 6QUW_A_GCP 6QUX_A_GCP 6T5B_A_GDP<br>6T5U_B_GDP 6T5V_A_GDP 6TAM_A_GDP 6TAN_A_GDP 6USX_A_GDP 6USZ_A_GDP<br>6UT0_A_GDP 6V5L_A_GDP 6V65_C_GNP 6V6F_C_GNP 6VC8_A_GNP 6VJJ_A_GNP<br>6W4E_B_GSP 6W4F_B_GDP 6WGN_A_GNP 6XGU_A_GNP 6XGV_A_GNP 6XHA_A_GNP<br>6XHB_A_GNP 6YR8_A_GDP 6YXW_A_GDP 6ZL5_A_GDP 6ZLI_B_GCP 7A1W_A_GDP<br>7A1X_A_GDP 7A1Y_A_GDP 7A47_A_GDP 7ACA_A_GCP 7ACF_A_GCP 7ACH_A_GCP<br>7ACQ_B_GDP 7C40_A_GDP 7C41_A_GDP 7EYX_A_GDP 7F0W_A_GDP 7KFZ_C_GNP<br>7LC1_A_GNP 7LC2_A_GNP 7LGI_A_GDP 7NY8_A_GDP 7ROV_A_GCP 7RSC_A_GSP<br>7RSE_A_GSP |
| CLASS-II | 4EPT_A_0QW 4EPV_A_0QX 4EPW_A_0QV 4EPX_A_0QR 4EPY_A_0QY 5OCO_B_9RK<br>6CC9_B_EWS 6CCX_B_EWS 6F76_A_CVK 6FA2_A_D2W 6FA3_A_D1Z 6FA4_A_D1W<br>6GJ5_A_F0N 6GJ6_A_EZZ 6GJ7_A_F0B 6GJ8_A_F0K 6GOM_A_F6E 6GQT_B_F8N<br>6GQW_A_F8T 6GQX_A_F8K 6GQY_A_F8Q 6V5L_A_QPD 6ZL5_A_F0K 6ZLI_B_QME<br>7A1X_A_QWB 7ACA_A_R6W 7ACF_A_R6W 7ACH_A_R6W 7ACQ_B_R6W 7LGI_A_P2Z |
| CLASS-III | 4LUC_A_20G 4LV6_A_20H 4LYJ_A_21F 4M1T_B_21M 4M1W_B_21R 4M1Y_B_21S<br>4M21_B_21Y 4M22_B_22C 5F2E_A_5UT 5V6S_A_8YD 5V6V_A_8YA 5V71_A_8ZG<br>5V9L_A_91D 5V9O_A_91G 5V9U_A_91S 5VBM_A_92V 5YXZ_A_94C 5YY1_A_94F<br>6B0V_A_C8G 6B0Y_A_8ZG 6FA1_A_D2Z 6N2J_A_K9M 6N2K_A_K9J 6OIM_A_MOV<br>6P8W_A_O67 6P8X_A_O5V 6P8Y_A_O5Y 6P8Z_A_O5S 6PGO_A_OJ1 6PGP_A_OHY<br>6T5B_A_O7K 6T5U_B_MKW 6T5V_A_MKZ 6TAM_A_MZQ 6TAN_A_MZN 6USX_A_M1R<br>6USZ_A_QH4 6UT0_A_M1X 7A1W_A_QWK 7A1Y_A_QWH 7A47_A_QY5 |

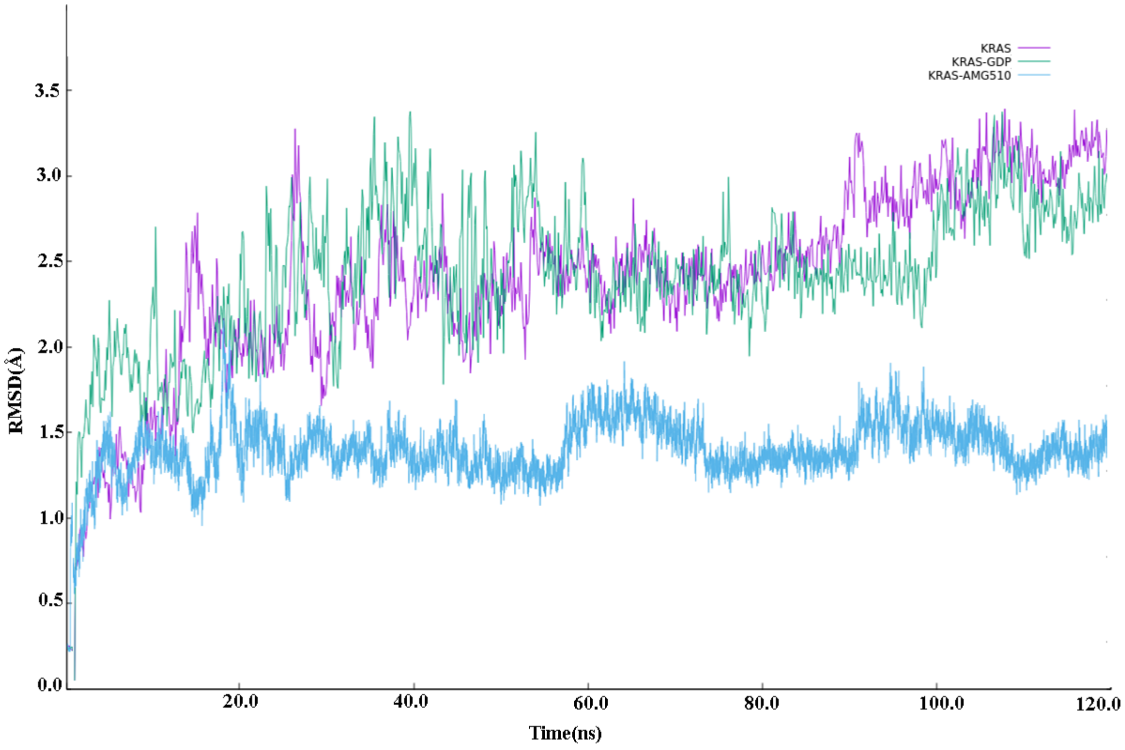


**Figure S1**. Ca-RMSDs of different trajectories for all three systems.

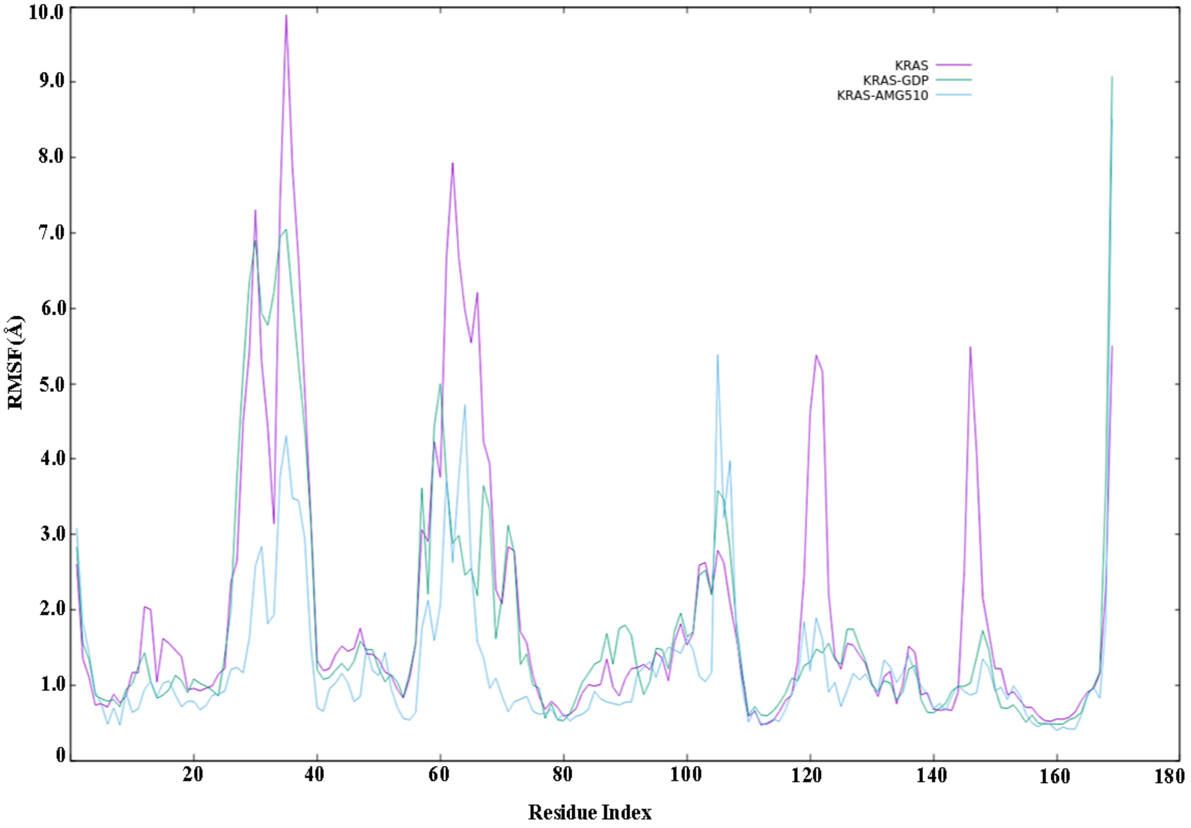


**Figure S2**. Ca-RMSFs of different trajectories for all three systems.
